# To mock or not: a comprehensive comparison of mock IP and DNA input for ChIP-seq

**DOI:** 10.1101/2019.12.17.880013

**Authors:** Jinrui Xu, Michelle M. Kudron, Alec Victorsen, Jiahao Gao, Haneen N. Ammouri, Fabio C. P. Navarro, Louis Gevirtzman, Robert H. Waterston, Kevin P. White, Valerie Reinke, Mark Gerstein

## Abstract

Chromatin immunoprecipitation (IP) followed by sequencing (ChIP-seq) is the gold standard to detect genome-wide DNA-protein binding. The binding sites of transcription factors facilitate many biological studies. Of emerging concern is the abundance of spurious sites in ChIP-seq, which are mainly caused by uneven genomic sonication and nonspecific interactions between chromatin and antibody. A “mock” IP is designed to correct for both factors, whereas a DNA input control corrects only for uneven sonication. However, a mock IP is more susceptible to technical noise than a DNA input, and empirically, these two controls perform similarly for ChIP-seq. Therefore, DNA input is currently being used almost exclusively. With a large dataset, we demonstrate that using a DNA input control results in a definable set of spurious sites, and their abundance is tightly associated with the intrinsic properties of the ChIP-seq sample. For example, compared to human cell lines, samples such as human tissues and whole worm and fly have more accessible genomes, and thus have more spurious sites. The large and varying abundance of spurious sites may impede comparative studies across multiple samples. In contrast, using a mock IP as control substantially removes these spurious sites, resulting in high-quality binding sites and facilitating their comparability across samples. Although outperformed by mock IP, DNA input is still informative and has unique advantages. Therefore, we have developed a method to use both controls in combination to further improve binding site detection.

## INTRODUCTION

ChIP-seq was developed to profile *in vivo* protein-DNA binding and histone modifications on a genomic scale (Barski et al. 2007; Johnson et al. 2007; Mikkelsen et al. 2007; Robertson et al. 2007). Compared to its predecessors, ChIP-seq has less noise and higher resolution (Schones and Zhao 2008; Ho et al. 2011), and thus is currently the standard technique to identify the binding sites of a transcription factor in the genome. ChIP-seq protocols typically begin with cross-linking DNA and its adjacent proteins using formaldehyde, followed by shearing DNA into small fragments by sonication. Next in the IP step, an antibody that binds specifically to the transcription factor (TF) of interest is used to enrich the TF-DNA complexes. Finally, the precipitated DNA fragments are sequenced and mapped back to a reference genome for binding site detection. The genomic regions with significantly more reads than controls are likely to be TF binding sites.

As with many high-throughput techniques, ChIP-seq is also susceptible to technical and biological biases (Park 2009; Kidder et al. 2011). In ChIP-seq, one bias arises during genome sonication, in which open chromatin regions are more easily sheared than other regions, and thus these open regions yield more protein-DNA complexes. Consequently, the IP step immunoprecipitates more complexes from the open chromatin regions, resulting in more sequencing reads. To correct this sonication bias, the fragmented genomes are divided into two portions. One portion goes through the IP step and then the sequencing step, whereas the other portion is sequenced directly to serve as input control. This direct sequencing result contains the shearing bias of sonication, and thus can be used to normalize the sequencing results from the IP protocol (Kharchenko et al. 2008).

In addition to sonication, uneven regulatory binding in the genome may result in bias during the IP step. For example, even without sonication bias, genomic regions with abundant DNA binding proteins tend to have more protein-DNA complexes. Although the antibody in IP binds specifically to its antigens, i.e. the target TFs, it can also bind nonspecifically to other proteins. Consequently, the antibody captures more protein-DNA complexes from genomic regions with abundant regulatory proteins. To control for this bias, a mock IP can be generated using the IP protocol, with the mock IP lacking specific antibody-antigen interactions. To this end, the mock IP either uses an antibody that cannot recognize the TF of interest, e.g., IgG, or the TF is not tagged with the epitope for the antibody used in the IP, e.g., GFP. Consequently, the mock IP control mimics only the nonspecific interactions in the IP. In addition to the nonspecific interactions, note that the mock IP also controls for sonication bias (Park 2009; Kidder et al. 2011; Landt et al. 2012). However, mock IP usually yields much less DNA material than DNA input and is thus more susceptible to technical noise (Kidder et al. 2011; Landt et al. 2012). Therefore, DNA input is recommended and used primarily in ChIP-seq (Park 2009; Meyer and Liu 2014). For example, in the ENCODE portal (Davis et al. 2018), almost all of the thousands of ChIP-seq data sets use DNA input as a control.

Increasing evidence suggests that spurious sites in ChIP-seq data may be substantial. Teytelman et al. and Park et al. find that TFs often appear to bind genomic regions that are counterintuitive to their function (Park et al. 2013; Teytelman et al. 2013). For example, TUP1 is recognized as a repressor of gene expression, but ChIP-seq still identifies its binding sites in the promoters of expressed genes (Park et al. 2013). Moreover, Jain et al., observed that when ChIP-seq was performed in a knockout background for a targeted TF, ~3,000 binding sites from the mutant fly embryos were still detected (Jain et al. 2015). The unexpected sites from these studies suggest the existence of abundant spurious sites. These potentially spurious sites tend to appear in highly transcribed genomic regions (Teytelman et al. 2013; Jain et al. 2015). Since DNA input controls are used in these examples, the potentially spurious sites are likely due to nonspecific interactions between the antibodies and other DNA-binding proteins or DNA fragments.

While these studies suggest the existence of spurious sites, interpretations of these results remain indefinite for two reasons. First, TF functionality is an ambiguous indicator of TF binding. Again, considering the TUP1 example - although generally considered as a repressor, TUP1 is also observed to activate genes (Zhang and Guarente 1994; Conlan et al. 1999). On the other hand, TF binding may not necessarily indicate biological function. Therefore, even though TUP1 usually acts as a repressor, when found in the promoters of transcribed genes, it may not be exerting any repressive functions. As a result, the binding of TUP1 detected around expressed genes may not be spurious. Secondly, the spurious binding sites in the aforementioned results were not predicted using the current standards and robust computational pipelines (Landt et al. 2012), and thus the numbers of spurious sites were sensitive to parameter settings.

Determining the abundance of spurious sites in ChIP-seq data is extremely important as the data are being widely used in numerous biological and medical studies. To this end, we generated and collected a large number of ChIP-seq datasets that have both mock IP and DNA input controls. We designed computational experiments that use these controls to estimate the abundance of spurious sites across different samples. Moreover, we proposed and validated that many spurious binding sites are potentially due to the intrinsic properties of samples. Such potential spurious sites can be removed using mock IP controls, but not using DNA input controls. Despite this result, our analyses indicate that DNA input controls are still informative for ChIP-seq. Therefore, we developed a novel method to utilize both mock IP and DNA input controls for improved binding site detection. This new tool can be used to tease apart biological binding sites from spurious ones to capture more accurate binding profiles of TFs.

## RESULTS

### ChIP-seq with multiple controls to illustrate the formation of spurious sites

#### Experimental setting of ChIP-seq data in use

Human ChIP-seq data were acquired from the ENCODE portal (Consortium 2012; Davis et al. 2018). The data were generated from six different cell line samples. Each sample has comparable DNA input (*d*) and mock IP (*m*) as controls for the IP experiments (*i*), as shown in figure 1A. In the IP experiment, an antibody specific to the target TF is used, whereas the mock IP uses an IgG antibody that does not specifically interact with any DNA binding proteins. In contrast, the ChIP-seq data of worm and fly are generated from whole organisms at the embryo, L4 and young adult stages in worm and the embryo, L3 larva and prepupae stages in fly. As shown in figure 1A, each of the worm and fly TFs has an IP experiment (*i*) and its DNA input control (*d*) generated by our modERN consortium (Kudron et al. 2018). In addition, for each of the stages, we produced a mock IP control (*m’*) as shown in figure 1B. We also generated DNA input controls (*d’*) corresponding to these mock IP controls (fig. 1B and Methods).

**Figure 1.**
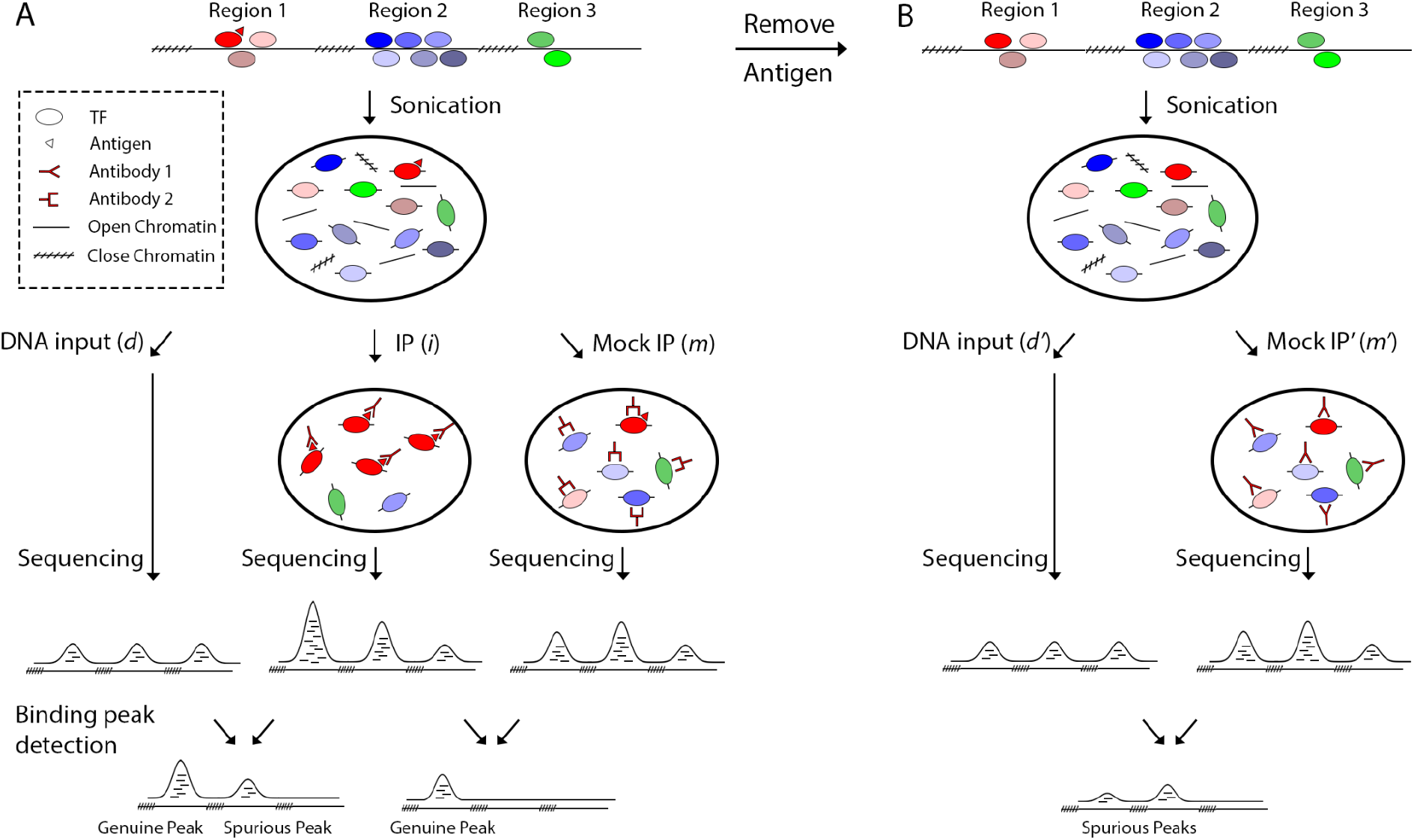
Illustration of the ChIP-seq protocols in use and the generation of spurious sites. The ChIP-seq protocol (A) can consist of IP, DNA input and mock IP experiments. For simplicity, the three open chromatin regions are assumed to be equally sensitive to sonication, and thus have three similar peaks of reads in the DNA input sequencing result. In IP, the peak of reads at region 1 is mainly due to the antibody specifically binding to the antigen (triangle) of the target TF (in red). The peaks at region 2 and 3 are due to nonspecific interactions between the antibody and regulatory proteins at the regions. In mock IP, to avoid the specific antigen-antibody reactions, we use another antibody, e.g. IgG, which does not bind specifically to any DNA binding proteins in the sample. Therefore, the resultant three peaks of reads are due to nonspecific interactions between the IgG and other DNA binding proteins. In this hypothetical example, a peak caller compares the three peaks of reads respectively from the IP and the DNA input, and then identifies binding peaks at region 1 and 2. Since there is no target TF binding at region 2, the detected binding peak is spurious due to strong nonspecific interactions at region 2. Using the mock IP as control, the peak caller identifies only the genuine binding peak at region 1. For worm and fly samples, due to the use of a GFP tag (see Methods), we can remove the antigen to avoid the specific reactions (B). Therefore, the mock IP for a worm or fly sample uses the same antibody (GFP antibody) as its IP. Because there is no antigen present in the sample for mock IP, the peaks of reads observed are also due to nonspecific interactions. A DNA input control is also generated for the worm or fly sample. The peak caller identifies binding peaks from the mock IP using the DNA input as control. However, due to lack of specific interactions, those binding peaks are all spurious.

#### Potential mechanism of spurious site generation due to nonspecific interactions

We propose a potential mechanism on how the nonspecific interactions between an antibody and regulatory proteins cause spurious sites when only DNA input controls are used. This mechanism is illustrated using a hypothetical example in Figure 1. For this purpose, we create three open chromatin regions and let them be equally sensitive to sonication. As a result, these regions have peaks of sequencing reads with similar heights in the DNA input control (*d*, fig. 1A). We let region 1 contain regulatory proteins as well as the target TF of the antibody. Regions 2 and 3 have no target TFs but only other regulatory proteins (fig. 1A). Therefore, in the IP experiment (*i*, fig. 1A), the peak of reads at region 1 is mainly due to the specific binding of the antibody to the target TF, whereas the peaks at regions 2 and 3 are purely due to nonspecific binding. Because we let region 2 have many more regulatory proteins than region 3 (fig1A), more complexes of regulatory proteins and DNA are generated from region 2. Therefore, even with nonspecific binding to the complexes and same sonication between region 2 and region 3, the antibody enriches more DNA fragments from region 2 than from region 3

With sufficient regulatory protein binding, the peak of reads at region 2 in the IP (*i*) can be higher than its counterpart in the DNA input (*d*), as we specified in figure 1. We postulate that this event may be further enhanced by physical and chemical factors at the molecular level. For example, the antibody used in the IP likely prefers to interact with the regulatory protein-DNA complexes from open chromatin rather than the histone-DNA complexes from closed chromatin. This preference may be due to the fact that regulatory proteins more likely resemble the target of the antibody than histones. Moreover, the histone-DNA complexes tend to carry no charge, which may further reduce binding to the antibody. Consequently, the antibody likely enriches more DNA fragments from open chromatin than from closed chromatin. This preference renders the peak of reads at region 2 in the IP even higher than that in the DNA input, taking the respective closed chromatin regions as reference. Due to the higher peak of reads at region 2 in IP than in DNA input, using the DNA input as a control for the IP results in a spurious binding peak at region 2.

This proposed mechanism of generating spurious sites predicts that between genomes, the ones with more abundant open chromatin and/or more highly expressed genes have a larger number of spurious binding sites. Moreover, this proposed mechanism also indicates that within a genome, the spurious sites tend to be associated with highly expressed genes, which recruit many regulatory proteins for transcription. This prediction is consistent with other recent observations (Park et al. 2013; Teytelman et al. 2013; Jain et al. 2015). However, spurious sites due to nonspecific interactions are expected to be removed when mock IP controls are used. As illustrated in figure 1, because the mock IP control (*m*) captures the nonspecific binding between the antibody and other regulatory proteins (Landt et al. 2012; Flensburg et al. 2014), the resultant peak of reads at region 2 in the mock IP is as high as the corresponding peak in the IP. In figure 1, region 3 has only a few regulatory binding proteins, and thus using either DNA input or mock IP control results in no spurious binding peak.

### Spurious binding sites across various samples

The abundance of spurious sites from nonspecific interactions between antibodies and regulatory proteins can be estimated by the sites detected from mock IP experiments compared to DNA input as control because mock IP experiments capture no specific interactions but only nonspecific ones (fig. 1B). Therefore, we used the standard ENCODE ChIP-seq pipeline to analyze the six pairs of mock IP (*m*) and DNA input (*d*) from the human cell lines, and the six pairs (*m’* and *d’*) from the worm and fly developmental stages. As a result, we observed that human cell lines on average have 9 spurious sites per 100 million base pairs (Mb) in genome, whereas this number increases to 551 and 3,931, respectively, for worm and fly samples. According to the mechanism we proposed, these numbers of spurious sites are expected to be correlated with the transcriptome activity and the genome accessibility of the samples.

In order to measure transcriptome activity, we used the RNA-seq data from the ENCODE portal (Davis et al. 2018) and our published data (Gerstein et al. 2014). However, the RPKMs from RNA-seq indicate the relative transcription levels of the genes within a sample, rendering across-sample comparisons impossible. For example, a cell type with all genes highly expressed has the same RPKMs as another cell type with all genes lowly expressed. To this end, we assume that the most highly transcribed genes of different eukaryote cell types have similar transcription activities, which are the limit of the transcription machinery. In a sample containing multiple cell types, the genes with the highest RPKMs are very likely the most transcribed genes in the majority cells. Taken together, we normalized the RPKMs of coding genes in each sample so that the five most highly expressed genes from the different samples had the same average, and then used the median of the normalized coding gene expression to indicate the transcriptome activity of the sample.

Note that for a sample with multiple cell types, the high median after normalization may be also because the many cell types express quite different genes from each other in the genome, indicating that on average a large fraction of the genome in the sample are actively transcribed. We also expect many spurious sites from such genomes. As a result, we used the medians of the normalized transcriptomes to roughly suggest their activities. These activities of the different samples are indeed highly correlated with the numbers of spurious sites per 100 Mb identified from the samples (Spearman’s *ρ* = 0.89, *P* < 9.2e-5). With linear regression, the transcriptome activity accounts for a large fraction of the variance in spurious site abundance (*r*^2^ = 0.92, *P* < 6.5e-7; fig. 2A). These results suggest that the large number of spurious sites seen in fly are due at least in part to its high transcriptome activity compared to worm and human cell lines.

**Figure 2.**
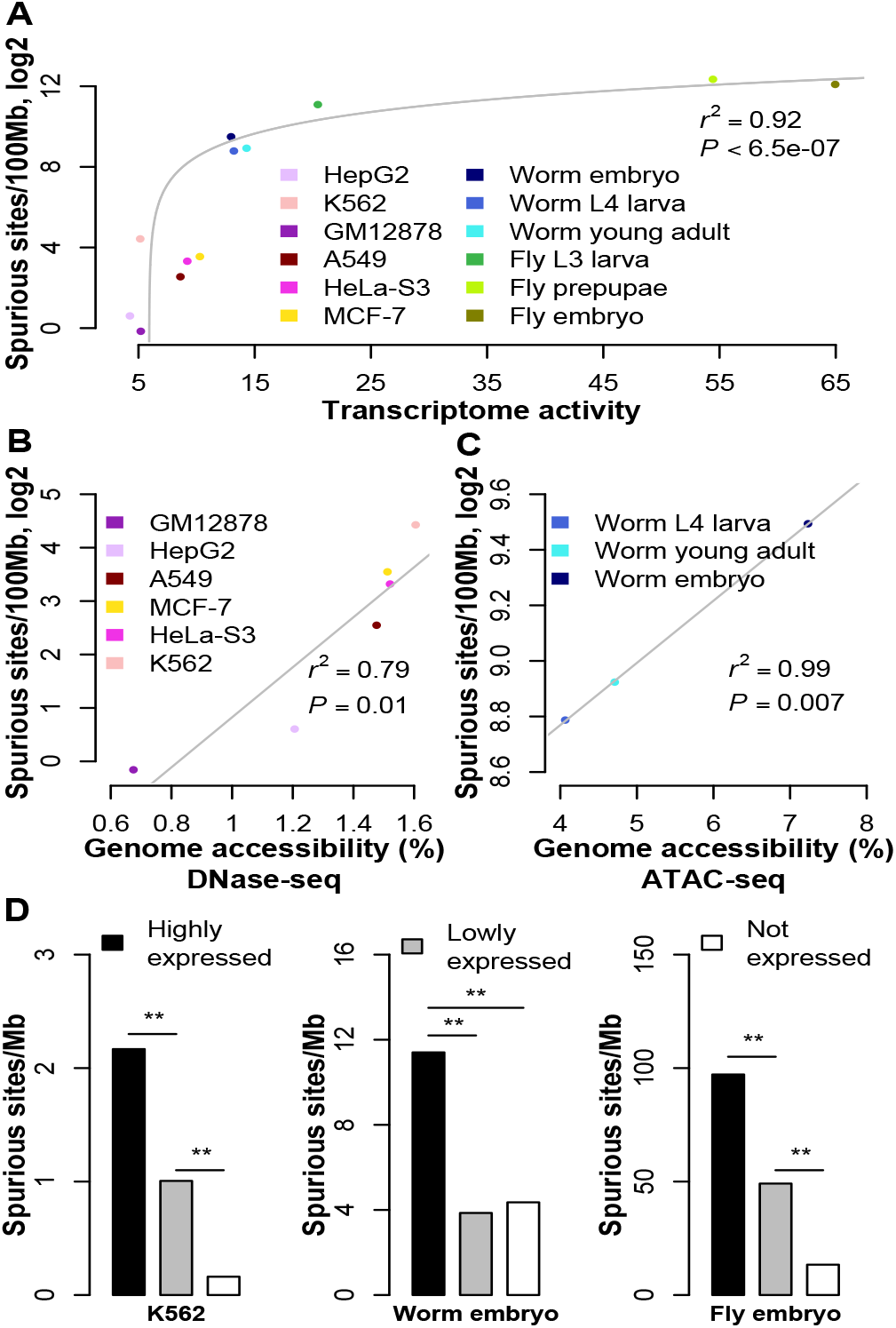
Associations between spurious site abundance and genome activities. The number of spurious sites per 100 million base pairs (Mb) in genome is linearly regressed with the transcriptome activity. For display purposes, the number of spurious sites is transformed into log space (A). The number of spurious binding sites is linearly regressed with genomic accessibility respectively for human samples (B) and worm samples (C). The number of spurious sites per 1 Mb is respectively calculated for promoter regions of highly expressed, lowly expressed and unexpressed genes in K562, worm embryo and fly embryo (D).

In addition, we tested the correlation between genome accessibility and spurious site abundance. The genome accessibilities of the six human cell lines were calculated from the DNase-seq data generated by the ENCODE consortium and processed uniformly by the ENCODE pipeline (Davis et al. 2018). The genome accessibility also explains a substantial fraction of the variance in spurious site abundance (*r*^2^ = 0.79, *P* = 0.01; fig. 2B). As expected, given their high growth rates, the five cancer cell lines have higher genome accessibilities than GM12878 cell line, and thus have more spurious binding sites (fig. 2B). For worm samples, Daugherty et al. generated ATAC-seq data and determined accessible regions from the data (Daugherty et al. 2017). The linear regression of these data again confirms the strong influence of genome accessibility on spurious site abundance as predicted (*r*^2^ = 0.99, *P* = 0.007; fig. 2C). Note that we paired the genome accessibility of L3 with the spurious site abundance of L4, due to lack of ATAC-seq data at the L4 stage. This mismatch renders the observed strong correlation between genome accessibility and spurious site abundance more conservative.

Our proposed mechanism of spurious site generation predicts that highly expressed genes tend to enrich more spurious sites than lowly expressed genes. To test this, we first classified coding genes as expressed and unexpressed (< 1 RPKM). The expressed genes were further split evenly into highly and lowly expressed genes. Gene-associated genomic regions were defined as 2kb regions both upstream and downstream of the transcription starting site(s). Within each of the three groups, overlapping associated regions were merged to avoid redundancy. Genomic regions that fell into multiple groups were reclassified into the highest expression group. For each of the three expression groups, the number of spurious sites whose summits lay within its genomic region was normalized by the total length of the region. For all the samples, we consistently observed that the promoter regions of highly expressed genes expression enriched with more spurious sites than other promoter regions (fig. 2D & S1). The GM12878, HepG2 and HeLa-S3 samples are excluded from this analysis due to insufficient spurious sites.

The human mock IP controls utilized IgG antibodies, while the fly and worm mock IP controls used the same GFP antibody. If the IgG antibody had a much higher specificity, i.e. less nonspecific interactions, than the GFP antibody, this higher specificity would result in the observed fewer spurious sites in the human cell lines than in the worm and fly samples. Although this difference in specificity is unlikely because both antibodies have been subject to quality control, we still tested this possibility by generating a mock IP in fly embryos using the IgG antibody. As expected, using IgG antibody leads to 6,059 spurious sites, which is similar to the 6,110 sites using the GFP antibody. This result supports that the difference in spurious site generation among human cell lines, worm and fly is not due to antibody specificity. In addition, the worm and fly samples used very similar ChIP-seq protocols and used the same library and sequencing protocols. Taken together, these results support that the observed differences in spurious site abundance across the species are due to the biological properties of the samples.

### Removing spurious binding sites from ChIP-seq using mock IP

Many spurious sites presumably due to the nonspecific interactions have been identified from mock IP using DNA input as control and these spurious sites may persist in IP experiments when DNA input controls are used. However, using mock IP as control may remove the spurious sites that persist in the IP (fig. 1A). To test this, we used the IPs of 113 ChIP-seq datasets across six human cell lines and 499 ChIP-seq datasets from the three stages each in worm and fly. These ChIP-seq data have matched mock IP and DNA input controls. Consistent with our prediction, using mock IP controls for binding site prediction results in fewer binding sites than using DNA input controls (fig. 3A). The reduction is marginal for human cell lines, but is substantial for worm and fly samples. As expected, this reduction is highly correlated with the ratio between the numbers of the spurious sites detected from mock IP and the total sites detected from IP, respectively using DNA input as control (fig. S2A). Expectedly, compared to the sites detected using the mock IP control, the sites obtained with the DNA input control are indeed more similar to the spurious sites (fig. S2B). Taken together, these results support the proposition that many spurious sites persist in IP experiments when DNA input controls are used, and mock IP controls remove a large number of such spurious sites.

**Figure 3.**
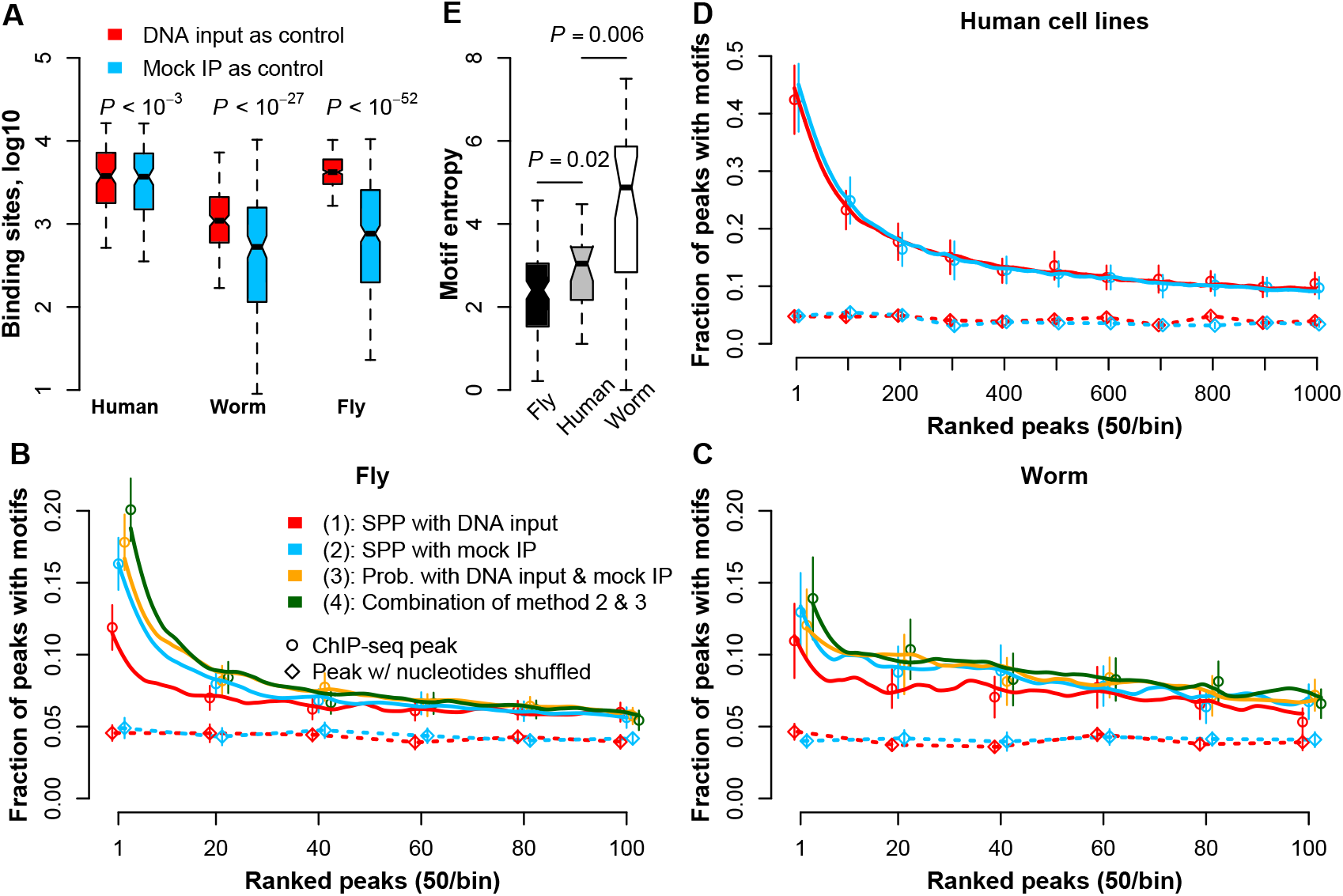
Reducing spurious sites from IP experiments using mock IP as controls. Compared to DNA input controls, the mock IP controls lead to fewer binding sites detected from IP experiments (A). And the binding peaks using mock IP controls also have higher motif enrichments in the fly samples (B), worm samples (C) and human cell lines (D). This higher enrichment is robust to different peak calling methods, e.g. SPP and our probability score. TF binding motifs of fly, human and worm have different entropies, depending on binding promiscuity and/or techincal ambiguity in determining the motifs (E).

### Motif enrichment in binding sites predicted using mock IP

Compared to using DNA input control, the mock IP control removes many potentially spurious sites. Therefore, the binding peaks detected using mock IP as control are expected to have higher motif enrichments than those using DNA input. To test this prediction, we examined 191 TFs in human, worm and fly with known binding motifs discovered by *in vitro* experiments such as protein-binding microarray (PBM) (Weirauch et al. 2014). As predicted, mock IP control substantially outperforms DNA input control in motif enrichment for fly and worm binding peaks (fig. 3B&C). The improvements are 37% and 18%, respectively, in the top 50 peaks ranked by the SPP signal (Kharchenko et al. 2008). As for human cell lines, the spurious site abundances are quite low, and thus the improvement in motif enrichment using mock IP as control is also small (*P* = 0.01; fig. 3D). To examine any GC-content effect, we shuffled the nucleotides of the predicted binding peaks to scramble motifs, but maintained the GC content of the motifs. We observed no difference in motif enrichment between using mock IP and DNA input controls (fig. 3B, C&D). This indicates that the observed improvements are not due to GC content differences between the sets of binding peaks in comparison.

Even using the mock IP control to normalize the IP samples, the predicted binding peaks of worm and fly TFs still have lower motif enrichments than those of human TFs (fig. 3B, C&D). However, we postulate that this comparison between TFs of different species is not informative for two reasons. First, the human, fly and worm motifs are generated using different techniques such as bacterial one-hybrid (B1H), systematic evolution of ligand by exponential enrichment (SELEX) and PBM (Tuerk and Gold 1990; Bulyk et al. 1999; Mukherjee et al. 2004; Meng et al. 2005). These high-throughput experiments may have quite different accuracies. Obviously, low accuracy diminishes the actual motif enrichment in binding peaks. Second, it may be that some TFs are more permissive than others and bind to a large number of various DNA sequences. As a result, for permissive TFs, even spurious sites tend to have binding motifs enriched, rendering the motif enrichment comparison between TFs not informative. Note that the existence of a motif does not necessarily indicate a TF binding event, which usually requires favorable chromatin states.

Both low motif accuracy and high TF promiscuity increase motif entropies, but the former artificially reduces motif enrichments in binding peaks, whereas the latter may artificially increase motif enrichments. We calculated the entropies of the motifs in worm, fly and human (fig. 3E; see Methods). A high entropy indicates that the DNA sequences of a motif are very diverse. Compared to fly motifs, the human ones have higher entropies (fig. 3E), and their enrichments are higher even in the peaks with low SPP binding signals, which are more likely to be spurious peaks (fig. 3D). Therefore, these results together suggest that the high entropies of human motifs may be in part due to TF promiscuity. This high promiscuity in human can be attributed to the lower constraint imposed by the smaller effective population size of human (~10^4^) (Rogers and Harpending 1992; Takahata 1993; Sherry et al. 1994; Erlich et al. 1996; Tenesa et al. 2007), compared to that of fly (~10^6^) (Hawks et al. 2000). As for worm, its effective population size (~10^4^) is comparable to that of human (Sivasundar and Hey 2003; Barriere and Felix 2005). However, compared to the human motifs, the worm motifs have much higher entropies (fig. 3E), but lower enrichments in the peaks with low SPP binding signals (fig. 3C&D). These results suggest that the extremely high entropy of worm is likely due to low motif accuracy. Taken together, TF promiscuity and motif accuracy may contribute to the different motif enrichments of the binding peaks among human, worm and fly.

### Predicting potential spurious site abundance in human tissues

We observed that spurious site abundance increases with sample complexity, i.e. from homogeneous human cell lines to worm and fly whole organisms. This increase is presumably because in each cell type, only certain genomic regions are open chromatin; however, aggregating multiple cell types creates more open chromatin, and thus more spurious sites. Consistent with this supposition, the human tissue/organ samples in the ENCODE portal have substantially more accessible chromatin than human cell lines/primary cells (fig. 4A). Unfortunately, there is a lack of comparable DNA input and mock IP controls for human tissues in the ENCODE portal, rendering direct estimation of spurious site abundance impossible. Instead, we used genome accessibilities of human tissues/organs to estimate their spurious site numbers. Extrapolating from the regression (fig. 2B), the median of spurious sites in human tissues and organs is 9,819, which is about 10-fold larger than the 849 predicted for all the available human cell lines and primary cells (fig. 4B). These rough estimations for human organs are especially important because ChIP-seq data on organs are widely used for studies of human diseases.

**Figure 4.**
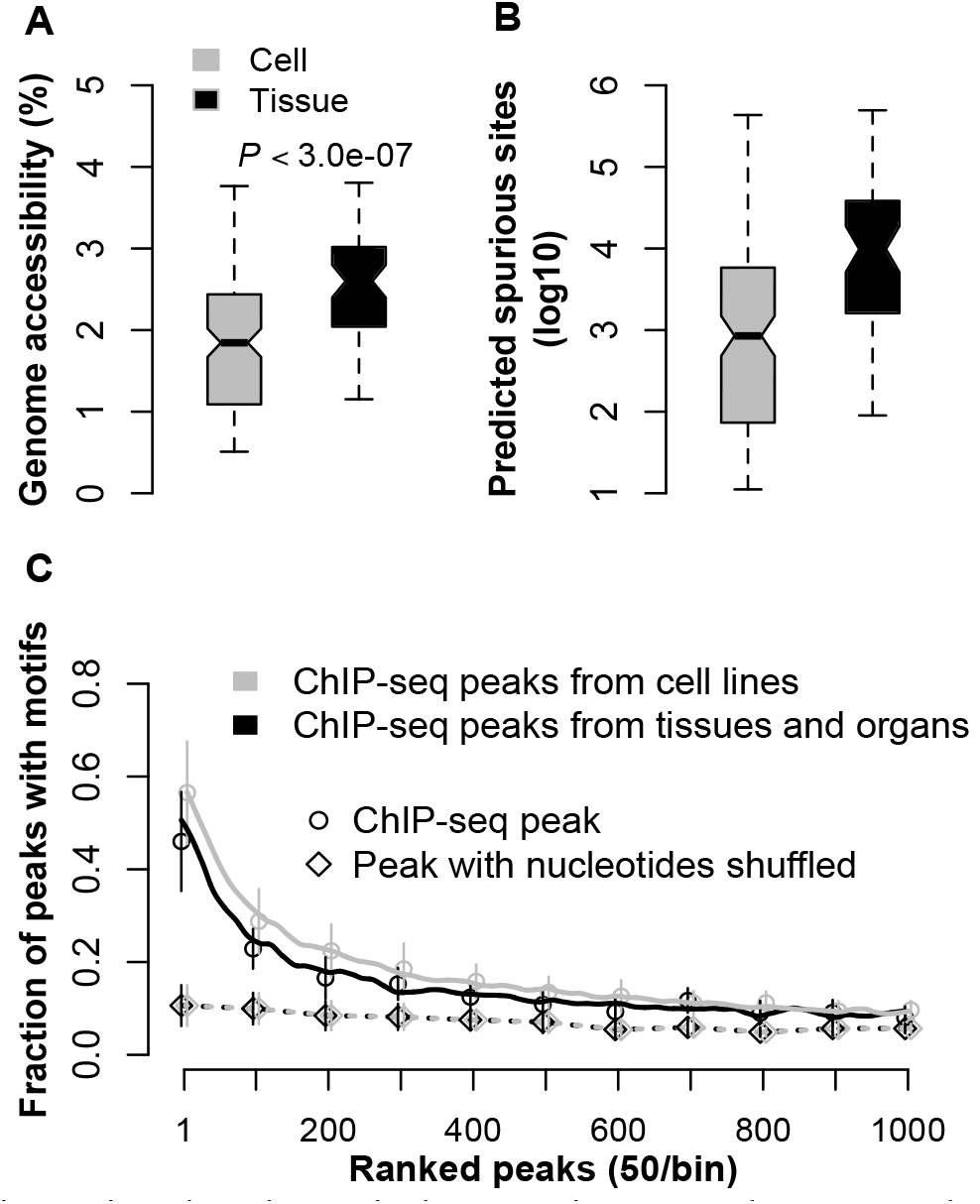
Inferring spurious site abundance in human tissues and organs. The human tissues and organs have much higher genome accessibilities than the human cell lines and primary cells (A). From the regression between genome accessibility and the number of spurious sites, the spurious site abudances are predicted from genome accessibilities for cell lines, primary cells, tissues and organs (B). Note that the number of spurious sites is defined as the sites detected from mock IP using DNA input as control. With more spurious sites in tissues and organs, their binding peaks detected from IP experiments using DNA input controls enrich less motifs than those of cell lines and primary cells (C).

The high spurious site abundance predicted in tissues/organs indicates low motif enrichment. As we discussed above, the motifs of different TFs are likely incomparable. Therefore, we focused on seven human TFs, i.e. RXRA, EGR1, SP1, MAX, GABPA, YY1 and HNF4A in the ENCODE portal. Each of the TFs has ChIP-seq data generated in both a human cell line/primary cell and a human tissue/organ. These TFs also have binding motifs determined by *in vitro* experiments in the Cis-bp database. With DNA input as control, the binding sites of these TFs in tissues indeed enrich fewer motifs than those in the cell lines (fig. 4C). This reduction is 17% for the top 50 binding peaks ranked by SPP binding signal. These results confirm the higher spurious site abundance in tissues/organs we predicted. Note that we used only genome accessibility for spurious site prediction, because adding transcriptome activity as another variable does not significantly improve the regression in figure 2B (*P* = 0.75), probably because transcriptome activity is merely an indirect measure of regulatory protein binding in the genome, compared to genome accessibility.

### Prevalence of potentially spurious sites in ChIP-seq using DNA input as control

We have detected spurious sites in mock IP experiments with DNA input controls. However, they do not necessarily all appear as spurious because of the interactions between antibodies and their target TFs in the IP experiments. Such specific interactions in IP may deplete non-specific interactions that cause spurious sites. Due to this depletion, the IPs are expected to have less spurious sites than the mock IP experiments. Particularly, the locus predicted as a spurious site in mock IP may also have strong binding of target TFs, and thus is identified as a *bona fide* site in IP. However, estimating the prevalence of persisting spurious sites in IP experiments is extremely difficult, if not possible, because the *bona fide* sites are unknown. To have a rough estimation, for each TF we first detect binding sites from its IP experiment, respectively with DNA input and mock IP as control, and then take the sites using mock IP as control as *bona fide* sites approximately. In the sites detected using DNA input control, the ones not overlapped with the approximate *bona fide* sites are the potentially spurious sites that remain.

This rough estimation demonstrates that on average a large fraction of predicted binding sites is potentially spurious in worm (60%) and fly (88%) samples, whereas this fraction is 10% in human cell lines (fig. 5A). This trend is consistent with the spurious site abundances observed across the samples. In addition to this sample-specific effect, we also found that canonically activating TFs annotated by the Gene Ontology (GO) database (Carbon et al. 2009) tend to have more binding sites detected by mock IP as compared to repressive TFs (fig. 5B); thus, activating TFs also have fewer spurious sites even when DNA input controls are used (fig. 5C). This trend is statistically significant for fly embryos, which have 115 assayed TFs annotated by GO (fig. 5C). In contrast, the trend is not significant for the samples of other developmental stages, presumably due to the small numbers of available TFs at those other stages (from 19 to 28 TFs). Note that the prevalence of the spurious sites depends on the accuracy of the reference sites taken as *bona fide*, which is largely unknown.

**Figure 5.**
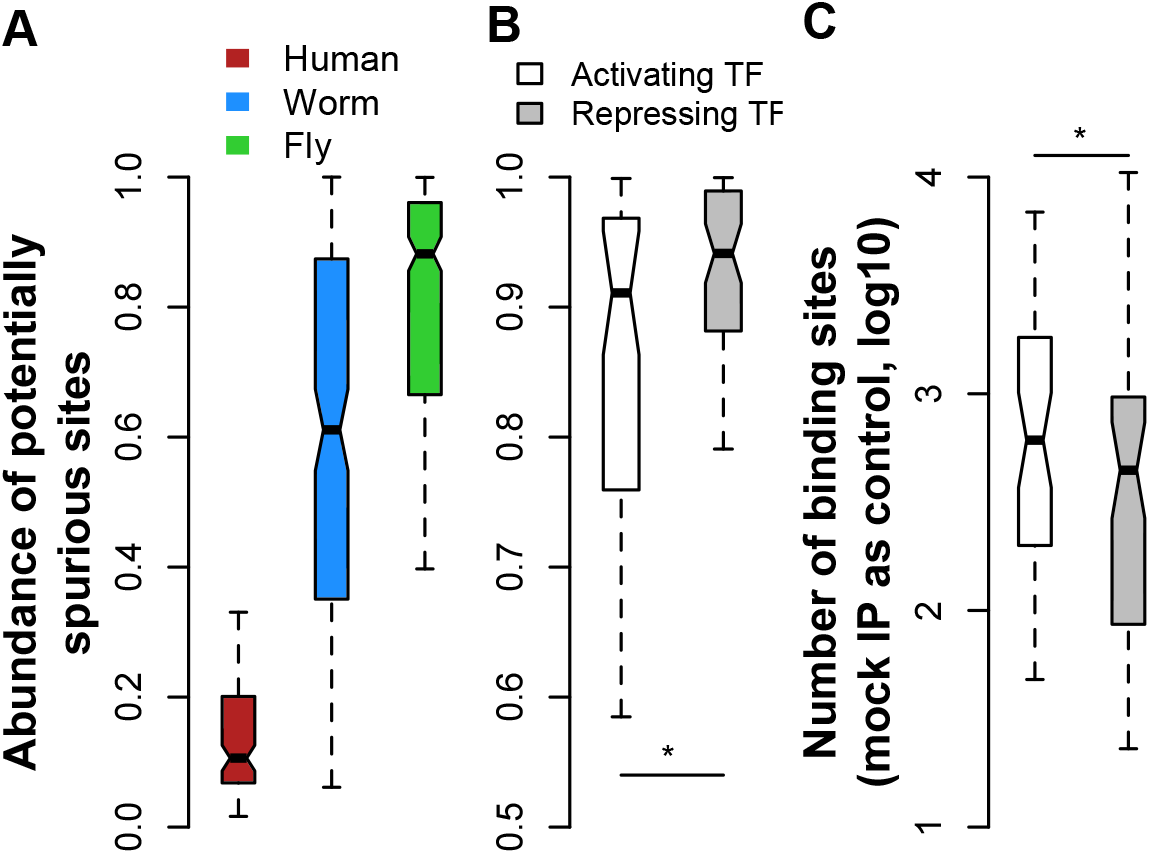
Spurious site prevalence is both sample specific (A) and TF specific (B, C). A large fraction of binding sites detected using DNA input as controls are potentially spurious in worm and fly samples, but not in the human cell lines (A). The activating TFs in fly embryo have more binding sites detected using mock IP controls than the repressing TFs (B). The activating TFs also have fewer spurious sites than the repressing TFs (C).

### Combining mock IP and DNA input to predict TF binding sites

Although the mock IP control removes spurious sites more efficiently than the DNA input control, the mock IP control usually acquires much less DNA material from a sample, and thus has a large variation in library preparation and sequencing. As a result, the mock IP controls are more prone to technical noise than DNA input controls. Therefore, we next tested whether combining the mock IP and DNA input controls together may further improve binding site quality. In addition, different scoring metrics have been developed for TF binding site detection. For example, the signal score in SPP depends directly on the enrichment of sequencing reads in the IP, compared to its control. Another metric, such as MACS, calculates the statistical significance of the read enrichment (Zhang et al. 2008). Combining these two scoring metrics may also improve binding site detection. To this end, we developed a simple framework that takes advantage of using both mock IP and DNA input controls as well as multiple scoring metrics in order to better detect binding sites. We focused on the ChIP-seq data of worm and fly TFs. Each of the TFs has an IP (*i*) and its DNA input control (*d*) as shown in figure 1A. The TF also has a mock IP (*m’*) with its own corresponding DNA input (*d’*) as shown in figure 1B.

First, we developed a Bayesian model to integrate the IP (*i*) and mock IP (*m’*) as well as their respective DNA input experiments (*d* and *d’*). These experiments were scaled to the same sequencing depth. For each TF, we used SPP to identify the peak regions in the genome using the IP (*i*) and the mock IP (*m’*). For each peak region, *n_i_*, *n_d_, n_m′_*, and *n_d′_* are the numbers of reads in the four experiments uniquely mappable to the region. The probability for the region being a binding site is indicated by *P*(*θ*_1_ > 0.5, *θ*_1_ > *θ*_2_), where *θ*_1_ = *n_i_*/(*n_i_* + *n_d_*) and *θ*_2_ = *n_m′_*/(*n_m′_* + *n_d′_*). We assume *n_i_* following a binomial distribution, i.e. *n_i_*~Bin(*n*_1_, *θ*_1_, where *θ*_1_~*U*(0,1) = Beta(1,1) and *n*_1_ is the number of total reads from the region in *i* and *d*. Because Beta(1,1) is an uninformative conjugate prior of the Bin(*n*_1_, *θ*_1_, the posterior distribution of *θ*_1_ is *P*(*θ*_1_|*n_i_*, *n*_1_ = Beta(*θ*_1_|*n_i_* + 1, *n_d_* + 1). With the same assumptions, we have *P*(*θ*_2_|*n_m′_, n*_2_) = Beta(*θ*_2_|*n_m_*, + 1, *n_d_*, + 1), where *n*_2_ is the number of total reads from the region in *m’* and *d’*. With this setting, *P*(*θ*_1_ > 0.5, *θ*_1_ > *θ*_2_) can be expressed as in equation 1.

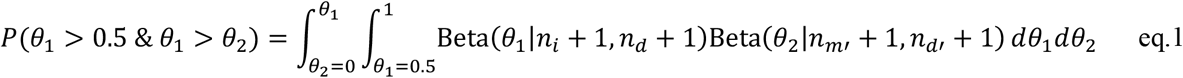

Due to lack of an analytical solution, we estimate the integral by simulation.

The higher probability indicates the genomic region is more likely to be a TF binding site. The genomic regions, as binding peaks, are ranked by this probability, and then passed to the tool of irreproducible discovery rate (IDR) (Li et al. 2011) in the ENCODE pipeline to determine binding sites. When DNA input is the only control, the probability is simply *P(J >* 0.5). With the same derivation, the probability of using only mock IP as control is also calculated as described in the Methods section. As expected, multiple controls substantially outperform respectively DNA input or mock IP alone (fig. S3). However, the probability with both controls performs similar to, but not always better than, the SPP score using only mock IP as control (fig. 3B&C). This observation is probably due to the fact that compared to the probability, the SPP score is more informative by considering not only the read enrichment but also the distribution of the reads at a genomic region.

Second, to take advantage of using both the SPP score and the probability calculated from both the mock IP and DNA input controls, we rank the peaks of a TF with the two scoring metrics respectively, resulting in two rankings for the one set of peaks. For each of the peaks, we sum its ranks in the two rankings and sort all the peaks again according to their summed ranks. This new ranking is then subject to IDR for binding site detection. This strategy is reasonable because IDR is a robust model that uses only the rank of each peak for binding site detection. This novel method increases motif enrichments by 21% and 8% in the top 50 binding sites, compared to the SPP method with mock IP control (fig. 3B&C). Currently, summing up the ranks of a peak implicitly assigns equal weights to the rankings by the two metrics. Using equal weights is appropriate in this case because the two metrics perform similarly in binding site detection (fig. 3B&C). In the case where one metric is substantially better than the other, the peak ranking by the more accurate metric can be multiplied by a large weight. The value of the weight reflects the credibility of one metric compared to the other. This strategy can be extended to multiple metrics with different weights since so many controls and scoring metrics have been generated during the past decade.

### Comparing the respective non-specific interactions of GFP and IgG antibodies

As aforementioned, using DNA input as controls, the mock IP experiments with the GFP antibody and the IgG antibody generate the similar numbers of spurious sites in fly embryos, suggesting that the two antibodies have the same amount of non-specific interactions. The interactions are non-specific because the interactants are not the specific antigens. However, the two antibodies may still prefer different interactants, resulting in different spurious sites. To this end, we compared the spurious sites of the IgG mock IP and GFP mock IP with DNA input as controls. Only 39% of the total spurious sites have overlaps in both of the mock IPs (fig. 6A). We generated another mock IP using GFP antibody for the fly embryo. Between the two GFP mock IPs, the overlapped spurious sites increase to 74% (fig. 6A). Due to the inconsistency between IgG and GFP antibodies and the fact that the IP experiments in ChIP-seq use GFP antibodies, the IgG mock IP is expected to have low efficiencies in removing spurious sites from the IPs. Indeed, using the IgG mock IP as control results in more sites and lower motif enrichments than using either of the two GFP mock IPs (fig. 6B&C). The two GFP mock IPs produce essentially the same numbers of binding sites and motif enrichments (fig. 6B&C).

**Figure 6.**
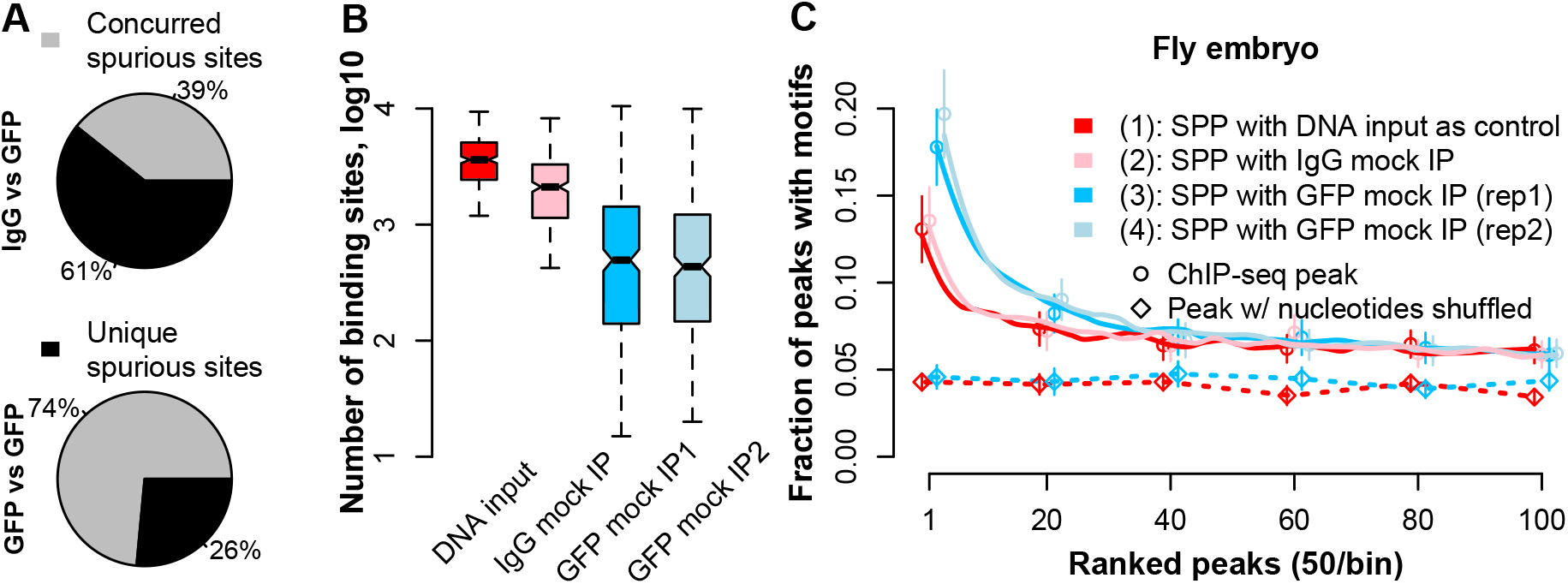
Comparing IgG mock IP and GFP mock IP in binding site detection for fly embryo. Two sets of spurious binding sites are detected from the IgG mock IP and the GFP mock IP, respectively using DNA input controls. A concurred spurious site in a set is the one that has an overlapping site in the other set. The fraction of concurred spurious sites between the IgG and GFP mock IPs is substantially smaller than that between the two GFP mock IPs (A). Using the IgG mock IP as control leads to much more sites from IP experiments than using the GFP mock IPs (B), and the resultant peaks have less motif enrichments (C).

## DISCUSSION

Our results indicate that many spurious sites exist in binding data generated by ChIP-seq experiments when DNA input is used as a control. Very roughly, we estimate that the fraction of spurious site ranges from 10% to 88% in our data sets (fig. 5), and our evidence indicates that the fraction depends on the genome accessibility and transcriptional activity of the sample (fig. 2). For example, samples such as tissues, organs, and whole organisms have much higher genome accessibility and transcriptional activity compared to cell lines and primary cells, and thus tend to have more spurious sites (fig. 2&4). Furthermore, with the same mechanism, cancer cell lines may have more spurious binding sites than normal cell lines (fig. 2B). Spurious binding sites tend to be enriched within the highly expressed genomic regions of a sample (fig. 2D). In essence, the uneven genome activity creates biases in IP experiments, and such biases cannot be fully controlled by DNA input experiments alone.

DNA input has been primarily used as controls in ChIP-seq experiments in various cell lines, tissues, and organs. The large fraction of spurious binding sites may impose various negative effects on applications of these ChIP-seq data. For example, including spurious binding sites with real TF binding sites, gene regulatory networks may artificially appear to be more connected than they are in reality. This ‘overconnected’ effect is expected to be more severe in tissues and organs than in cell lines. Moreover, the varying pervasiveness of spurious sites across samples reduces the comparability of ChIP-seq data. For instance, our results suggest that cancer cell lines may have more spurious sites than normal cell lines, rendering the TFs in cancer cell lines may artificially have more binding targets than in normal cell lines. In addition, the widely observed association between spurious sites and highly expressed genes further exacerbates the comparability of ChIP-seq data. Nevertheless, even with spurious sites, the sites predicted using DNA input controls still have substantial binding motif enrichments, which is especially true for the most confidently predicted sites. Moreover, our analyses suggest that strong TF binding depletes spurious sites. Intriguingly, for example, if the TFs in a cancer cell have higher expression levels than those in a normal cell, the binding of these TFs might be enhanced in the cancer cell. As a result, the predicted sites from cancer cells may have more *bona fide* sites and less spurious sites, compared to those from normal cells.

To address these issues, we used mock IP as controls to detect binding sites of many TFs in human cell lines, worm and fly. Because mock IP closely mimics the systematic biases caused by genome activity, the mock IP control reduces spurious sites substantially. This removal increases the quality of predicted binding sites. In the samples with many spurious sites, we found that mock IP control increases the motif enrichment of top binding sites by 18-37%, compared to DNA input control (fig. 3B&C). Besides the systematic bias, ChIP-seq, especially the mock IP, suffers from random technical noise. Therefore, we developed a novel framework that utilizes both mock IP and DNA input controls as well as multiple scoring metrics for binding site detections. This combination further improves binding site quality by 8-21% (fig. 3B&C). By reducing spurious sites, using mock IP controls increases the comparability of predicted binding sites across different samples. This novel method and the large set of binding sites we generated are valuable assets for ChIP-seq applications.

We have demonstrated that the mock IP control can remove spurious sites because it captures the systematic biases in ChIP-seq. In theory, the biases can be removed by optimizing many other factors in ChIP-seq, such as antibody quality, library preparation, the buffers used, fragment size selection, sequencing depth and cross-linking time, etc (Kidder et al. 2011; Gilfillan et al. 2012; Landt et al. 2012; Meyer and Liu 2014; Baranello et al. 2016). However, in practice, these multiple factors may be antagonistic, and may require independent optimization for different samples and TFs. Therefore, optimizing these factors for each sample and TF is not practical for determining the binding sites of a large number of TFs in various samples. This is particularly true for our modERN project which aims to detect the binding sites of all TFs in different developmental stages of both worm and fly. In contrast, incorporating a mock IP control requires much less effort, and improves binding site accuracy.

Our discoveries also provide potential insight for highly occupied targeted (HOT) regions, which have previously been identified in ChIP-seq data across many species (Moorman et al. 2006; Gerstein et al. 2010). A HOT region is a genomic region with binding sites of more TFs than expected. Intriguingly, some HOT regions function as developmental enhancers and have distinct regulatory signatures (Kvon et al. 2012). However, increasing evidence hints that some HOT regions may be artificial due to the false binding signals from ChIP-seq (Wreczycka et al. 2019). Our results confirm the existence of abundant spurious sites in ChIP-seq, and thus the potential influence on HOT regions. Quantitatively, spurious sites are expected to artificially inflate the numbers of TFs in some HOT regions. However, our results suggest that spurious sites are unlikely to influence HOT regions qualitatively because the existence of spurious sites in a genomic region indicates abundant regulatory protein binding in that region, and thus the region is likely to have more TF binding than expected.

The three species, human, worm, and fly, have similar numbers of coding genes as potential targets of TFs. However, using mock IP controls, TFs in human cell lines tend to have substantially more binding sites than TFs assayed in whole worm or fly, and the fly TFs have slightly more binding sites than the worm TFs (fig. 3A). These numbers of binding sites across the three species are proportional to their genome sizes. This proportion to the genome size may be due to the fact that the larger genome contains more motifs for TF binding. The more motifs in the larger genome may be favored by natural selection to attract TFs around the chromosomes, which increases the utility of TFs. Moreover, a larger genome may have more “TF reservoirs”, which are DNA sequences containing weak binding affinities to TFs and might be used to buffer the system and maintain an optimal amount of available TFs in the nucleus (Lin and Riggs 1975; MacQuarrie et al. 2011).

In summary, we provide evidence for a potential mechanism and a corrective approach to address the issue of spurious site abundance in ChIP-seq data. The abundance of spurious sites in a sample is strongly associated with its genome accessibility. Early ChIP-seq studies in humans have focused mainly on cell lines, which have short accessible regions, and thus small numbers of spurious sites. For these samples, using DNA input and mock IP controls perform similarly, which might have led to the widely believed notion that a DNA input control is sufficient for ChIP-seq. However, in samples such as tissues and whole organisms like worm and fly, the abundance of spurious sites is substantial and increases with overall genome accessibility. We have demonstrated that these spurious sites can be removed using mock IP controls, with the resulting binding sites becoming more accurate and comparable across samples. For further improvement, we developed a novel method that incorporates both DNA input and mock IP controls as well as scoring metrics for binding site detection. The enhanced binding site detection method will better capture the true binding sites of TFs to gain a better understanding of their roles in development and physiology.

## METHODS

### ENCODE pipeline for peak calling and binding site detection

The ChIP-seq data for each TF includes at least two IP experiments as biological replicates and a control. The high-quality reads of each set are uniquely aligned to a reference genome using BWA (Li and Durbin 2009) And the mapped reads of the two IP replicates are pooled together first, and then the pooled reads are randomly divided into two sets, which are pseudo-replicates. These biological replicates, pseudo-replicates, and the control set are used by the established ENCODE pipeline for peak calling and then binding site prediction (Landt et al. 2012). In the pipeline, SPP is used to detect TF binding peaks by comparing an IP experiment to its control experiment (Kharchenko et al. 2008). As a result, for each of the IP sets and pseudo-replicate sets, a list of TF binding peaks is detected and ranked by the SPP score. These multiple lists of ranked binding peaks are passed to the irreproducible discovery rate (IDR) tool to determine binding sites (Li et al. 2011). To estimate spurious sites, we replace IP experiments by mock IP experiments.

### ChIP-seq data of human, worm and fly from ENCODE and modERN

#### ChIP-seq data of human cell lines, tissues, and organs from ENCODE

We acquired ChIP-seq data of human TFs from the ENCODE portal (Davis et al. 2018) and focused on six cell lines, namely GM12878, K562, HepG2, A549, HeLa-S3, and MCF-7 because both mock IPs and DNA input controls are available for each of these cell lines. Each pair of the mock IP and DNA input experiments is assigned as controls to an IP experiment of the corresponding cell line. Across the six cell lines, we utilized 113 ChIP-seq data with an IP experiment, a DNA input and a mock IP paired with the input (Table S1). We excluded the ChIP-seq data for histone marks, polymerases and CTCF from our analyses. In addition, we found seven human TFs with ChIP-seq data and DNA input controls from both cell lines or primary cells, and tissue or organ samples (Table S2). These ChIP-seq data were used to compare the spurious site abundances between simple and complex samples.

#### ChIP-seq data of worm and fly from modERN

We used 317 and 182 ChIP-seq data generated by our modERN consortium for the whole organisms of worm and fly respectively (Tables S3&4). In detail, the worm ChIP-seq data are from developmental stages of embryo, L4 and young adult. As for fly, the ChIP-seq data are from embryo, W3L and WPP developmental stages. The modERN ChIP-seq protocol tags the target TF with the green fluorescent protein (GFP), generating a transgenic fly or worm. The same GFP antibody is used in both organisms during the IP process. A detailed protocol for strain generation and ChIP-seq is described in (Kudron et al. 2018). The ChIP-seq data for each TF consists of 2 replicate IP experiments along with a DNA input control. For each of the 3 developmental stages in worm and fly, we generated mock IP and DNA input control samples. The mock IP was performed using the GFP antibody in wild type animals that did not contain a tagged TF, thus the GFP antibody has no specific antigen in these samples (Table S5).

### Comparable transcriptome activities across human, worm and fly samples

The gene expression levels of the six human cell lines were measured by RNA-seq in the ENCODE portal (Table S6) (Davis et al. 2018). As for the developmental stages of worm and fly, the RNA-seq data were generated by (Gerstein et al. 2014). The fly and worm embryonic stage gene expression levels were measured across many time points. Therefore, we averaged the gene expression levels over the time points to estimate the gene expression level of the embryonic stages as a whole. We then focused on the coding genes of each species, because their annotations are more accurate across the three species. While RNA-seq measures relative gene expression within a sample, the gene expression values are not comparable across samples. To address this, we multiplied the gene expressions of each sample by a specific number so that the top 5 highly expressed coding genes of all the samples have the same average. The median of the scaled expressions of a sample was used to indicate the transcriptome activity of the sample.

### Genome accessibilities of human and worm samples

We used DNase-seq data and ATAC-seq data respectively to measure the accessibilities of human and worm samples. The DNase-seq data of the six cell lines were collected from the ENCODE portal (Davis et al. 2018) and were generated by the ENCODE consortium and processed uniformly by the DNase-seq pipeline of ENCODE to detect accessible regions in the genomes of the samples (Table S7). The total length of the accessible regions was used to indicate the genome accessibility of a human sample. As for worm, we used ATAC-seq data generated and processed uniformly by (Daugherty et al. 2017). They assayed worm samples at embryo, L3 and young adult stages. These stages match our ChIP-seq stages, except for the L4 stage. Therefore, we used the L3 ATAC-seq for the L4 stage ChIP-seq. This slight mismatch renders our hypothesis testing more conservative. Similar to the DNase-seq data, we used the total length of accessible regions to indicate the genome accessibility of a worm sample. We also acquired 60 DNase-seq datasets from human tissues and organs (Table S7). This number is 150 from all human cell lines and primary cells (Table S7). From their genome accessibilities, we predicted the numbers of spurious binding sites in the samples.

### Binding site detection using posterior probability as a scoring metric

For a given TF, we focus on the IP (*i*), DNA input (*d*) and mock IP (*m*) experiments. These experiments are all scaled to the same sequencing depth. With the DNA input control (*d*) for the IP (*i*), we use the SPP in the ENCODE pipeline to identify the binding peak regions in genome. For each peak region, *n_i_* and *n_d_* are the numbers of reads in the respective experiments mapped to the region. The likelihood for the region being a binding site is indicated by *P*(*θ* > 0.5), where *θ* = *n_i_*/(*n_i_* + *n_d_*). We assume *n_i_*~Bin(*n, θ*), where *θ*~Beta(1,1) and *n* is the number of total reads from the region in *i* and *d*. And thus the posterior distribution of *θ* is *P*(*θ*|*n_i_,n*) = Beta(*θ*|*n_i_* + 1, *n_d_* + 1). Instead of the DNA input (*d*), we also use the mock IP as control (*m*). The same model setting results in *P*(*θ′*|*n_i_*, *n′*) = Beta(*θ′*|*n_i_* + 1, *n_m_* + 1), where *n′* is the number of total reads from the region in *i* and *m*. These two probabilities are used respectively for binding site detections.

### Motif enrichment in TF binding regions

From the Cis-BP database (Weirauch et al. 2014), we collected the position weight matrix files (PWM) of motifs determined by *in vitro* methods such as systematic evolution of ligands by exponential enrichment (SELEX) (Tuerk and Gold 1990), protein-binding microarray (PBM) (Bulyk et al. 1999; Mukherjee et al. 2004) and bacterial one-hybrid (B1H) (Meng et al. 2005). Occasionally, some TFs have multiple PWMs, which are often determined in different research publications. For such a TF, we randomly selected one of the multiple motifs for analysis. For the 127 fly ChIP-seq data, the TFs have known motifs (Table S8). For worm and human respectively, the numbers are 29 and 44 (Table S8).

We used FIMO to search for motif hits *(P* < 10^-4^) in the reference genomes (Grant et al. 2011). For a binding site detected by the ChIP-seq pipeline, we define its core region as 100 bp around the summit as determined by SPP, and thus each binding site is considered as a 200 bp region. For binding sites of a TF, its motif enrichment is defined as the fraction of the binding sites containing the known TF motifs. Using this fraction renders motif enrichments comparable across different TFs and different species, whose binding sites may have quite different numbers of motifs. To generate a control for GC content, we divide a reference genome into 10 bp bins, and then shuffled the sequence within each bin. Such shuffling breaks the motifs, if any, in a binding site, while preserving the GC contents of the motifs.

### Motif entropy calculation

The entropy of a motif is calculated from its position weight matrix (PWM). Each element in the matrix is denoted as *P_k,j_*, which is the frequency of the nucleotide *k* at the *j*th position of the motif. The *k* represents one of the four nucleotides. Therefore, the entropy of a motif is calculated as in equation 2,

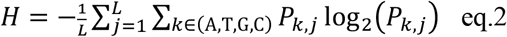

where *L* is the length of the motif.

## DATA ACCESS

All raw and processed sequencing data generated in this study have been submitted to the ENCODE portal. In case the data has not been released by the portal, they are also provided in the supporting information. The supplementary figures and tables are also in the supporting information.

## SUPPORT INFORMATION

***Supplementary figures***

http://archive2.gersteinlab.org/proj/MockOrNot/SupportInformation/Supplementary_Figures.docx

***Supplementary tables***

http://archive2.gersteinlab.org/proj/MockOrNot/SupportInformation/Supplementary_Tables.xlsx

***Supplementary data***

For the manuscript, we generated seven sets of mock IP data. These data have been submitted to the ENCODE portal (Table S5). The fastq files also can be accessed at our website: http://archive2.gersteinlab.org/proj/MockOrNot/Data/Raw/fastq/

In this manuscript, the TF binding sites in worm and fly whole organisms are detected respectively using DNA input, mock IP and combined controls.

These sites are deposited in http://archive2.gersteinlab.org/proj/MockOrNot/Data/Processed/

The corresponding directories are ChIP-seq_wDNAInput, ChIP-seq_wMockIP and ChIP-seq_Combo.

Particularly, for human cell lines, the TF binding sites are only detected using DNA input and mock IP controls, respectively.

These binding sites are in ChIP-seq_wDNAInput and ChIP-seq_wMockIP, respectively.

The other IP experiments and DNA input controls are downloaded from the ENCODE portal.

These experiments of human, worm and fly TFs are listed in the supplementary tables.

